## Supplementary Information_Figures for "PCLAF-DREAM Drives Alveolar Cell Plasticity for Lung Regeneration"

**List of Supplementary Materials**

**Supplementary Figures 1 to 19**
**Supplementary Tables 1 to 8**

**Supplementary Table 1. Marker genes of each cell type in mouse lung scRNA-seq data**
The list of marker genes for each cell type identity of mouse lung scRNA-seq data (GSE1412259), analyzed by the 'FindAllMarkers' function of Seurat.

**Supplementary Table 2. Marker genes of each cell type in human lung scRNA-seq**
The list of marker genes for each cell type identity of human lung scRNA-seq data (GSE135893), analyzed by the 'FindAllMarkers' function of Seurat.

**Supplementary Table 3. Marker genes of each cell type in mouse lung scRNA-seq from *Pclaf* WT and *Pclaf* KO mice at 7 dpi of bleomycin**
The list of marker genes for each cell type identity of *Pclaf* WT and KO mouse lung epithelial cells (collected from mice at 7 dpi of bleomycin instillation), analyzed by 'FindAllMarkers' function of Seurat.

**Supplementary Table 4. List of gene sets from of input into CLUE database**
The list of gene sets of input into the CLUE database. Gene sets of the PAPCs (*Pclaf* WT vs. *Pclaf* KO scRNA-seq shown in Figure 2). Gene sets of KP cells (*shPclaf* vs. *shControl [shCon]*) and H1792 cells (*shPCLAF* vs. *shControl [shCon]*) were identified from the bulk RNA-seq data (GSE136571 and GSE147305, respectively).

**Supplementary Table 5. List of drug candidates identified from the CLUE database**
This spreadsheet has the output of the CLUE database with each gene sets. Each output was derived from the input gene sets listed in Supplementary Table 4; gene sets of PAPCs (*Pclaf* WT vs. *Pclaf* KO), KP cells (*shPclaf* vs. *shControl [shCon]*) and H1792 cells (*shPCLAF* vs. *shControl [shCon]*). CLUE outputs (drug candidates) with scores higher than 1.5 were selected.

**Supplementary Table 6. Gene ontology of DREAM-target genes generated from the PANTHER database**
The list of gene ontology (GO) (molecular function) of DREAM-target genes analyzed by the PANTHER db [1]. The gene set of DREAM-target genes (968 genes; FISCHER_DREAM_TARGETS) was analyzed for the statistical overrepresentation test of GO molecular function complete.

**Supplementary Table 7. Antibody and primer information**
This spreadsheet has all information related to antibodies (manufacturers, catalog numbers, dilution rates, and antigen retrieval methods) and primers (sequences for cloning and mRNA quantification).

**Supplementary Table 8. Gene sets used for module score analysis**
The list of gene sets used for module score using with scRNA-seq dataset. List of genes in DREAM target genes, Sox9-based progenitor genes, MYC target genes, SMAD3 target genes from A549 cells, SMAD3 target genes from mouse embryonic cells, and SMAD3 target genes from human embryonic cells.

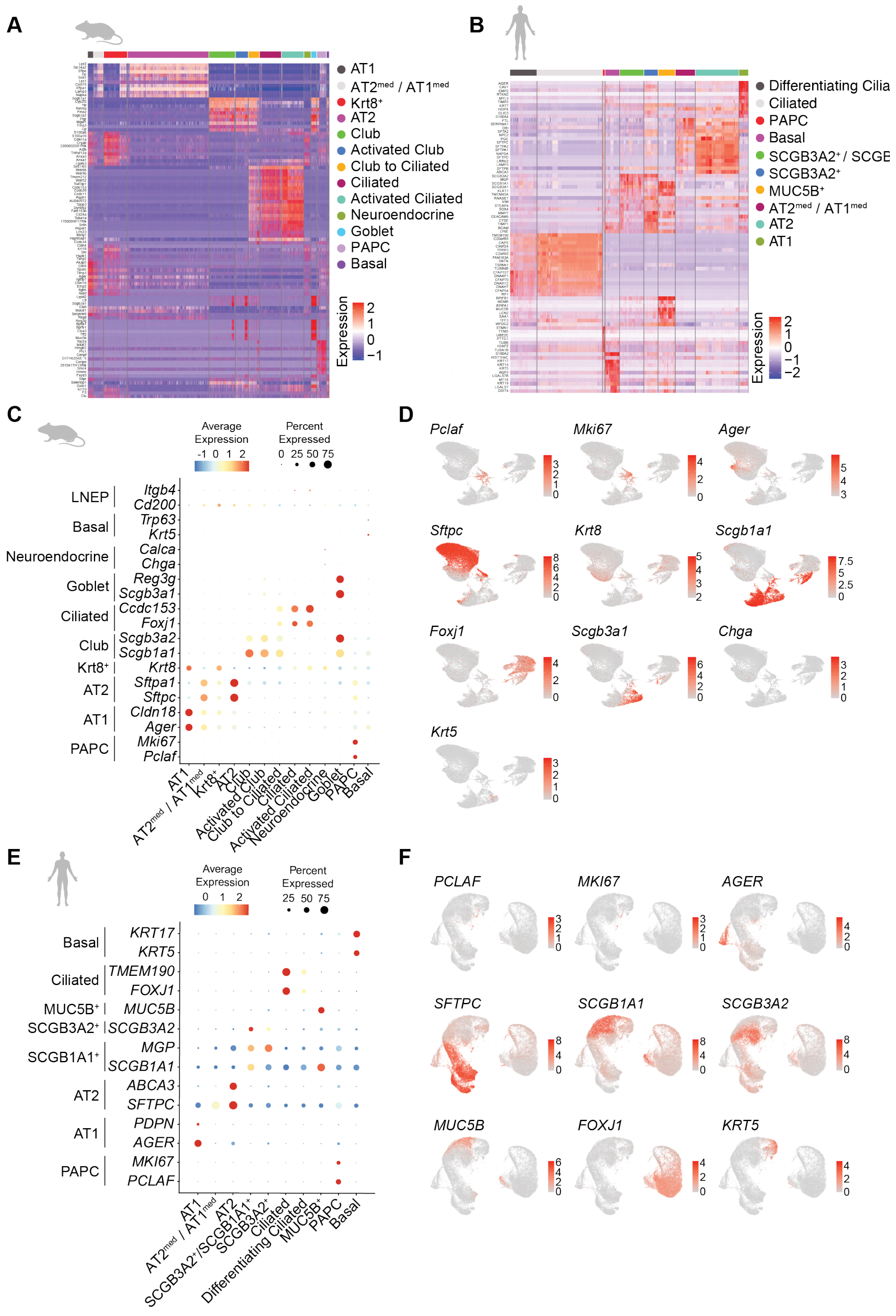


**Supplementary Fig. 1. Cell type annotation of human and mouse lung single-cell transcriptomics**

**A** Heatmap of gene expression of the top 10 genes in each cell type of mouse lung using data shown in Figure 1A. **B** Heatmap of gene expression of the top 10 genes in each cell type of human lung using data shown in Figure 1D. **C** Dot plots for mouse lung epithelial marker gene expression in each cell type. **D** Feature plots of mouse lung epithelial marker gene expression. **E** Dot plots for human lung epithelial marker gene expression in each cell type. **F** Feature plots of human lung epithelial marker gene expression.

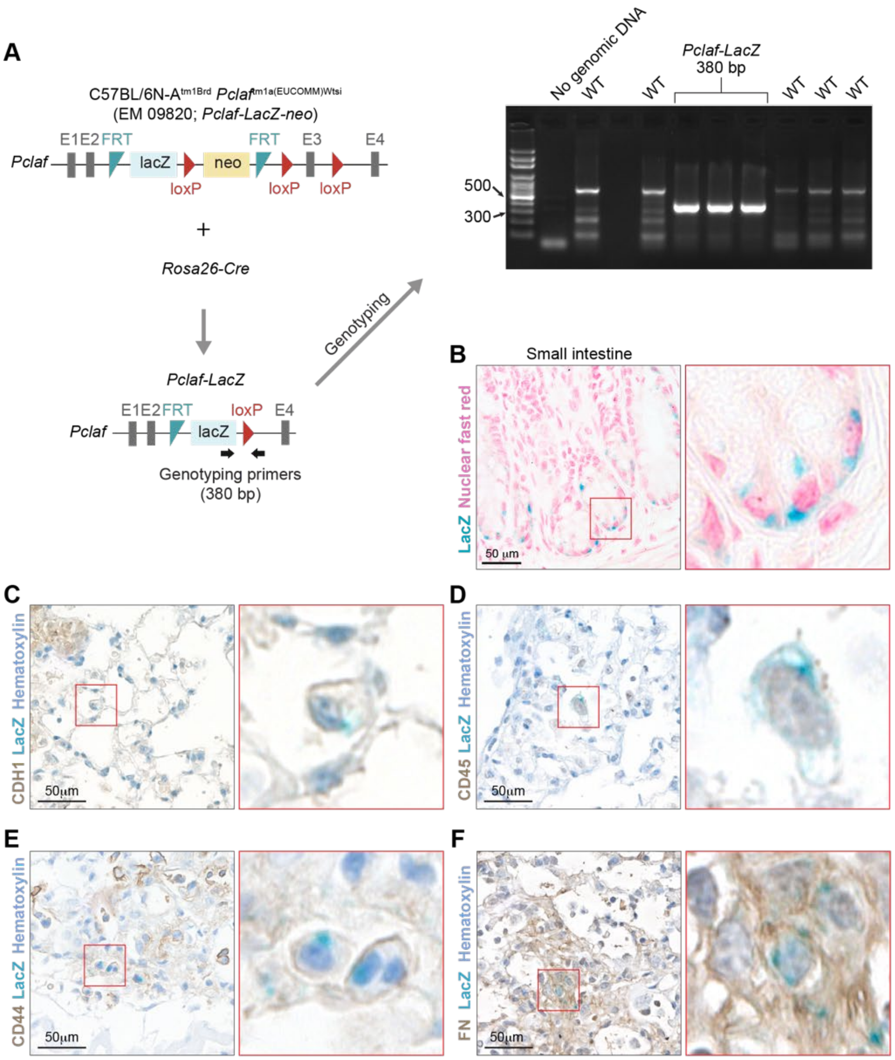


**Supplementary Fig. 2. Experimental scheme for generating *Pclaf-LacZ* mice**

**A** Experimental scheme for generating *Pclaf-LacZ* mice. C57BL/6N-A$^{tm1Brd}$ *Pclaf*$^{tm1a(EUCOMM)Wtsi}$/WtsiPh mice were bred with the *Rosa26-Cre* strain. The *Pclaf-LacZ* allele was confirmed by PCR-based genotyping. **B** Representative image of X-gal staining of the small intestine where PCLAF is specifically expressed in the crypt [2]. PCLAF-LacZ was expressed in the crypt base columnar cells. **C-F** *Pclaf-LacZ* mice were treated with bleomycin (1.4 U/kg, by intratracheal instillation). At 7 dpi, lungs were collected and processed for immunostaining: epithelial cell (**C**; CDH1), immune cell (**D**; CD45), or mesenchymal cell (**E**; CD44 and **F**; fibronectin (FN)) in combination with X-gal staining. The represented images are shown (n≥3).

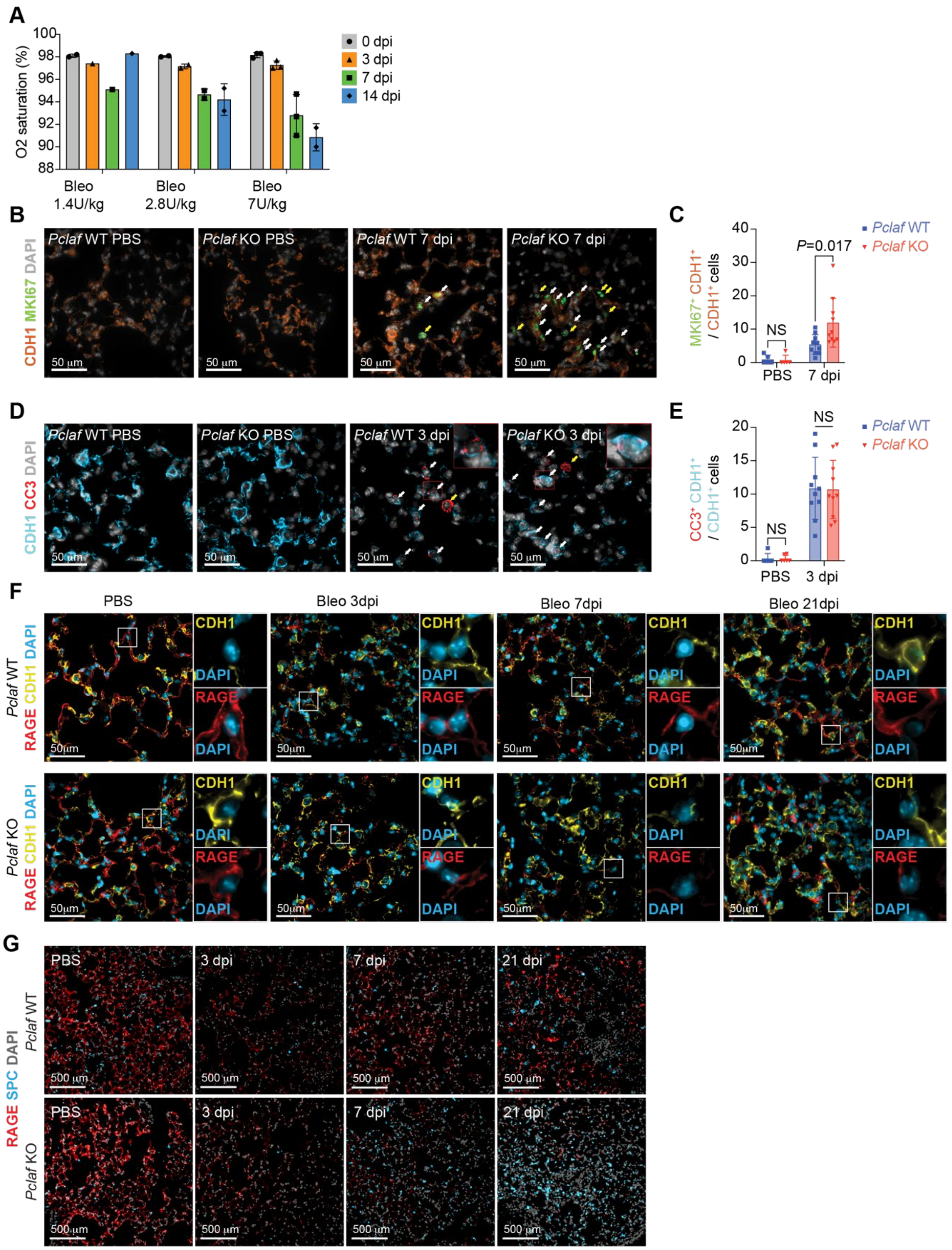

**Supplementary Fig. 3. *Pclaf* KO inhibits AT1 regeneration and promotes AT2 repopulation.**
**A** 8-week-old C57BL/6 mice were treated with bleomycin (1.4 U/kg; n=2, 2.8 U/kg; n=2, 7.0 U/kg; n=3; intratracheal instillation). The dynamics of $SpO_2$ levels were measured by pulse-oximetry at the indicated time points. One mouse instilled with bleomycin (1.4 U/kg) was dead at 1 dpi. Mouse instilled with 1.4 U/kg bleomycin showed recovered $SpO_2$ level back to as levels of at 0 dpi at 14 dpi, while mice with 2.8 U/kg or 7 U/kg of bleomycin showed inhibited $SpO_2$ level at 14 dpi. **B-G** Experimental scheme for the bleomycin-induced lung injury model. *Pclaf* WT and KO mice were treated with phosphate-buffered saline (PBS) (n=5 for *Pclaf* WT, n=5 for *Pclaf* KO) or bleomycin (1.4 U/kg; n=25 for *Pclaf* WT, n=24 for *Pclaf* KO) by intratracheal

instillation. Representative images of immunostaining for CDH1 and MKI67 at indicated time points (**B**). Quantification graph of $CDH1^{+}/MKI67^{+}$ cells / $CDH1^{+}$ cells (**C**). Representative images of immunostaining for CDH1 and cleaved caspase 3 (CC3) at indicated time points (**D**). Quantification graph of $CDH1^{+}/CC3^{+}$ cells / $CDH1^{+}$ cells (**E**). Representative images of immunostaining for RAGE and CDH1 at indicated time points (**F**). Representative images of immunostaining for RAGE and SPC at indicated time points (**G**). The represented images are shown (n≥3).

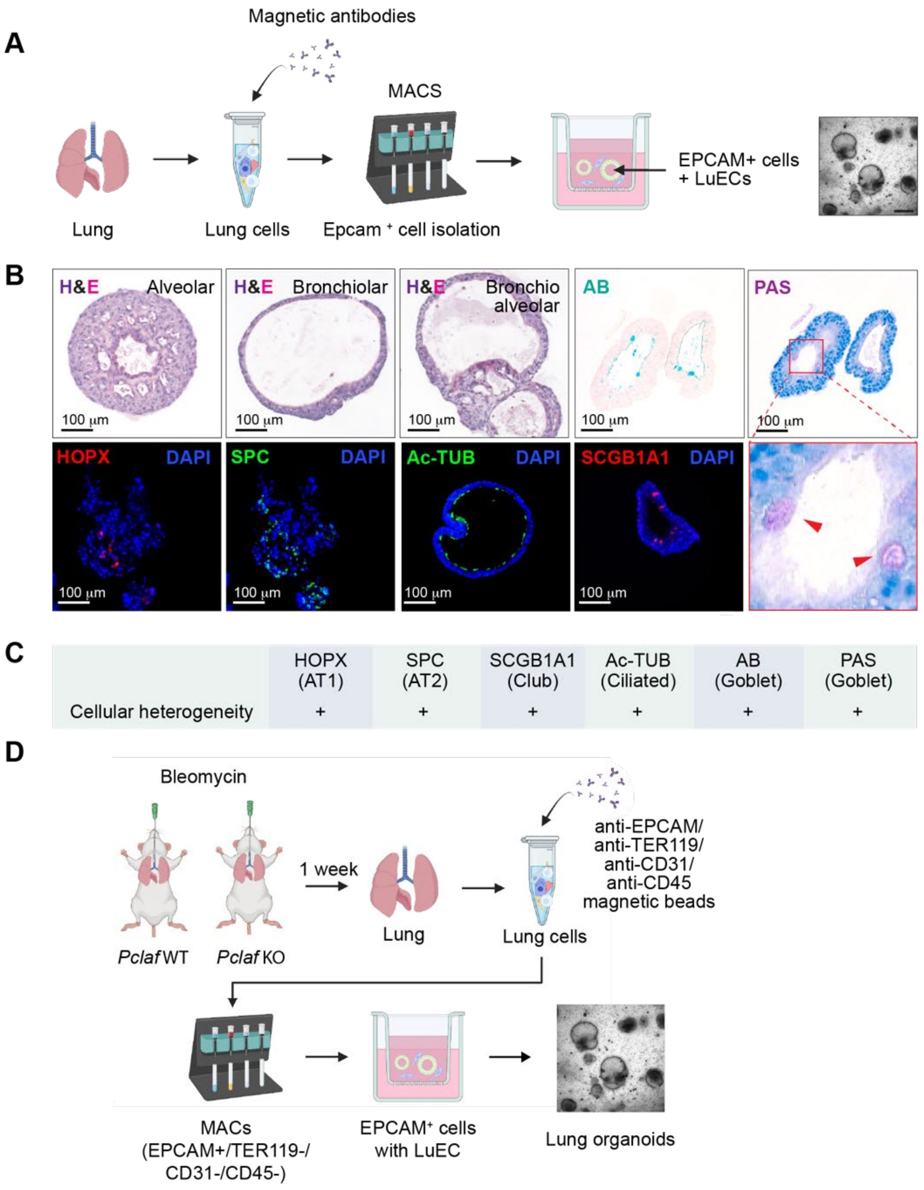


**Supplementary Fig. 4. Lung organoid culture system**

**A** Scheme of LO culture. The lung epithelial cells were isolated from WT mice by magnetic-activated cell sorting (MACS) and cultured with LuECs at a liquid-air interface to grow LOs. **B** Representative images of organoids for hematoxylin and eosin (H&E), alcian blue (AB), and periodic acid-Schiff (PAS) staining (*upper* panels), and immunostained for HOPX (AT1 cells), SPC (AT2 cells), acetyl-Tubulin (Ac-TUB; Ciliated cells), and SCGB1A1 (Club cells) of organoids with LuECs. **C** Table of positive cells using data shown in Supplementary Fig 4B. **D** Scheme of LO culture. The lung epithelial cells were isolated from bleomycin-treated lungs of *Pclaf* WT or *Pclaf* KO mice at 7 dpi by MACS. The lung epithelial cells ($TER119^-$/$CD31^-$/$CD45^-$/$EPCAM^+$) were cultured with lung endothelial cells ($CD31^+$) at a liquid-air interface to generate LOs. The represented images are shown (n≥3).

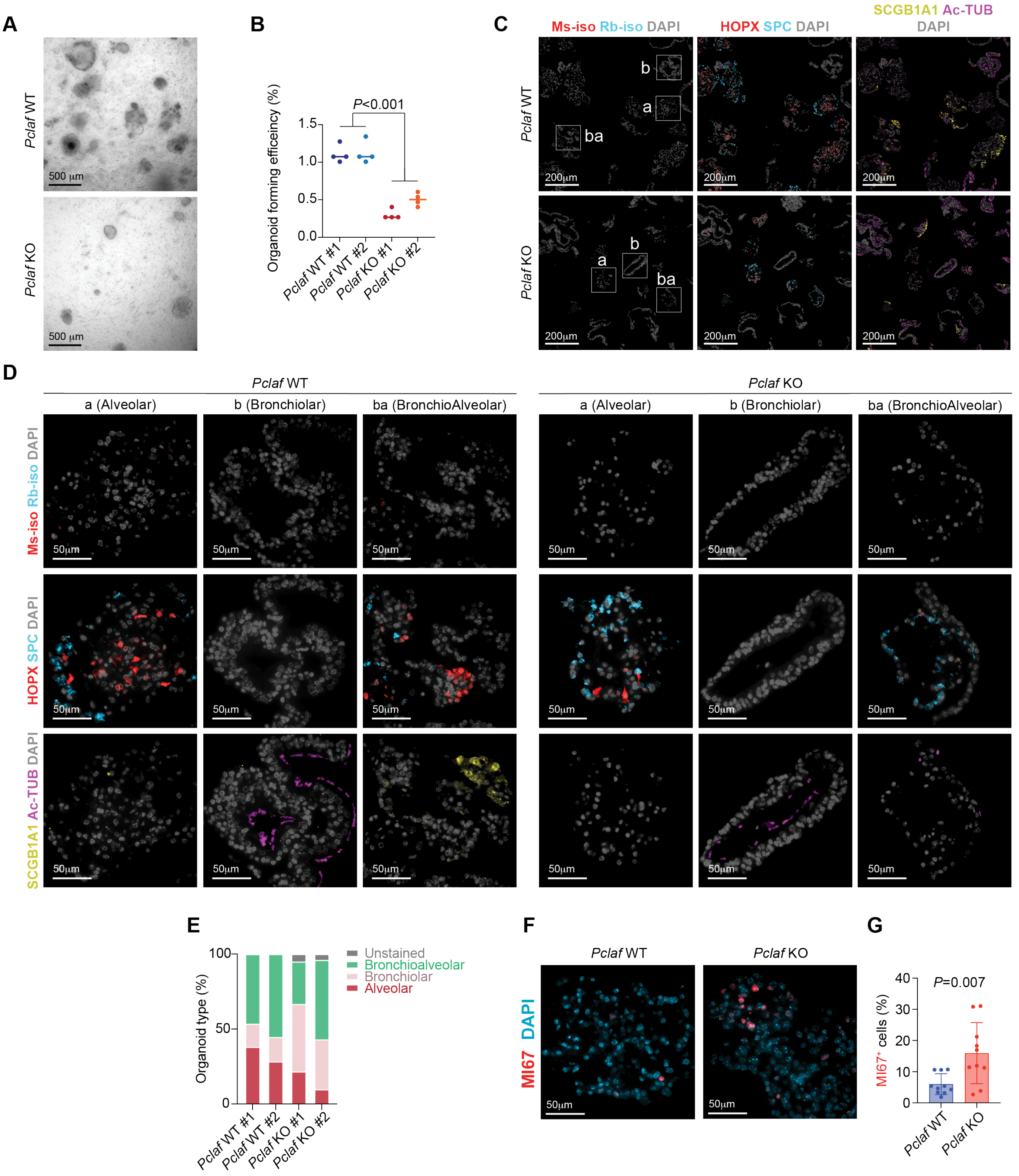


**Supplementary Fig. 5. *Pclaf* KO inhibits alveolar-type lung organoid formation.**
The lung epithelial cells were isolated from bleomycin-treated lungs of *Pclaf* WT or *Pclaf* KO mice at 7 dpi (bleomycin administration) by magnetic-activated cell sorting (MACS). The lung epithelial cells (TER119$^-$/CD31$^-$/CD45$^-$/EPCAM$^+$) were cultured with lung endothelial cells (CD31$^+$) at a liquid-air interface to generate LOs. **A** Bright-field z-stack images of LOs at day 12. **B** Quantification graph of lung organoid forming efficiency (OFE). **C** Representative images of LOs that were fluorescently immunostained for RAGE (AT1

cells), SPC (AT2 cells) Ac-TUB (Ciliated cells), and SCGB1A1 (Club cells). Isotype-controls (rabbit IgG for SPC and SCGB1A1, mouse IgG for HOPX and Ac-TUB) served as negative controls. Serial sections were used for isotype-control, HOPX and SPC, and Ac-TUB and SCGB1A1 staining. a; alveolar, b; bronchiolar, ba; bronchioalveolar. **D** High magnification images of each alveolar, bronchiolar, and bronchioalveolar organoids shown and indicated in Supplementary Figure 5C. **E** Quantification graph of organoid type. **F** Representative images of LOs that were fluorescently immunostained for MKI67. **G** Quantification graph of $MKI67^+$ cells / $DAPI^+$ cells. #1 and #2 indicate two independent experiment sets. Two-tailed Student's *t*-test; error bars: SD. The represented images and data are shown (n≥3).

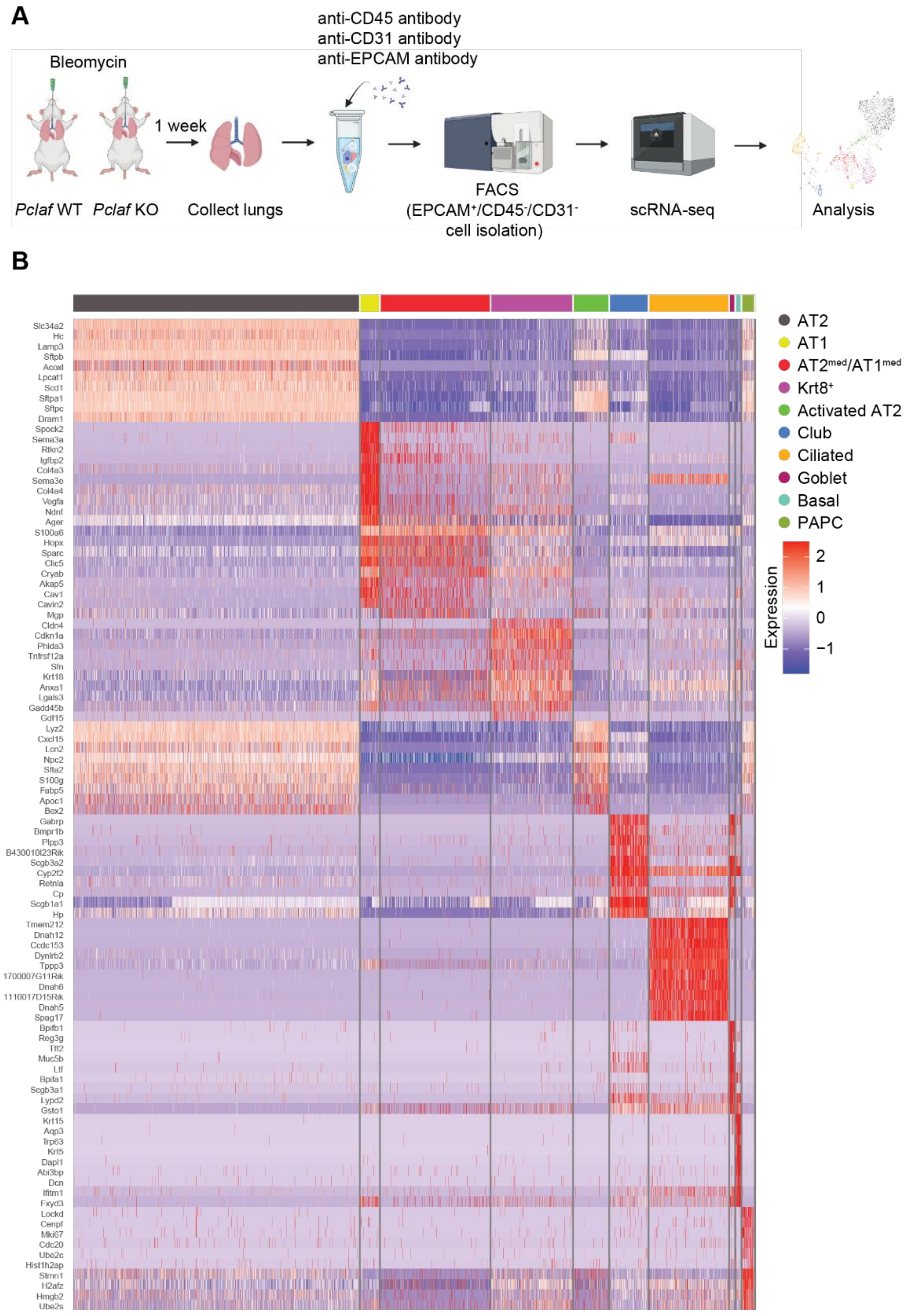


**Supplementary Fig. 6. Scheme of single-cell transcriptomics of *Pclaf* WT and KO mouse lung tissues**
**A** Scheme of scRNA-seq. Lung epithelial cells were isolated from *Pclaf* WT and KO mice (treated with bleomycin) by fluorescence-activated cell sorting (FACS). $CD31^{-}/CD45^{-}/EPCAM^{+}$ cells were used to generate sequencing data using the 10X genomics single-cell sequencing platform. **B** Heatmap of gene expression-based cell clusters of the top 10 genes in each cell type of mouse lung using data shown in Figure 3B.

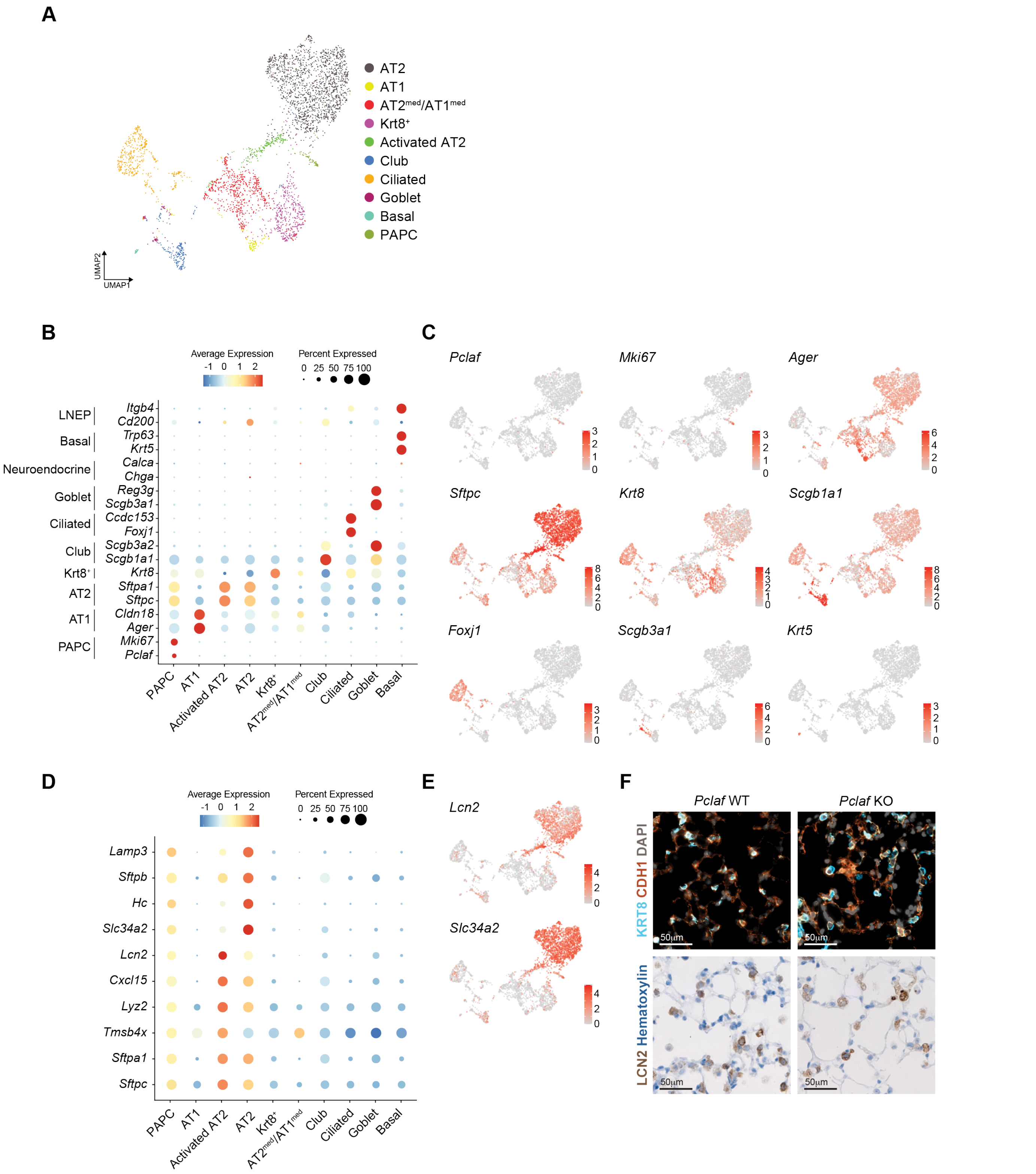


**Supplementary Fig. 7. Cell type annotation of mouse lung single-cell transcriptomics**

Analysis of the scRNA-seq dataset is shown in Figure 3B. **A** UMAP displays each cell cluster, colored by cell type. **B** Dot plots for mouse lung epithelial marker gene expression quantification of each cell type. **C** Feature plots of mouse lung epithelial marker gene expression. **D** Dot plots for genes specifically expressed in AT2 and activated AT2 cells. **E** Feature plots displaying the expression of *Lcn2* and *Slc34a2*. **F** Representative images of immunostaining for KRT8 and CDH1, or LCN2 using lung slides at 7 dpi as shown in Figure 2. The represented images and data are shown (n≥3).

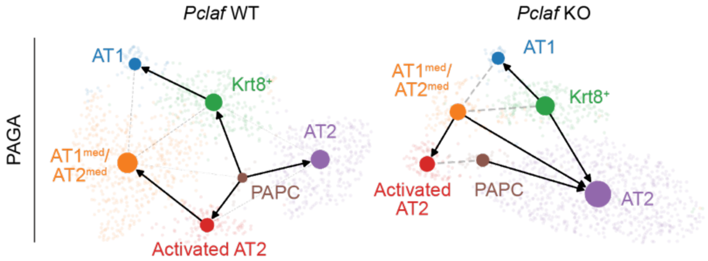


**Supplementary Fig. 8. *Pclaf* KO-impaired cell lineage trajectory from AT2 cells to AT1 cells**
Partition-based graph abstraction (PAGA) analysis using the results of RNA velocity analysis shown in Figure 3C.

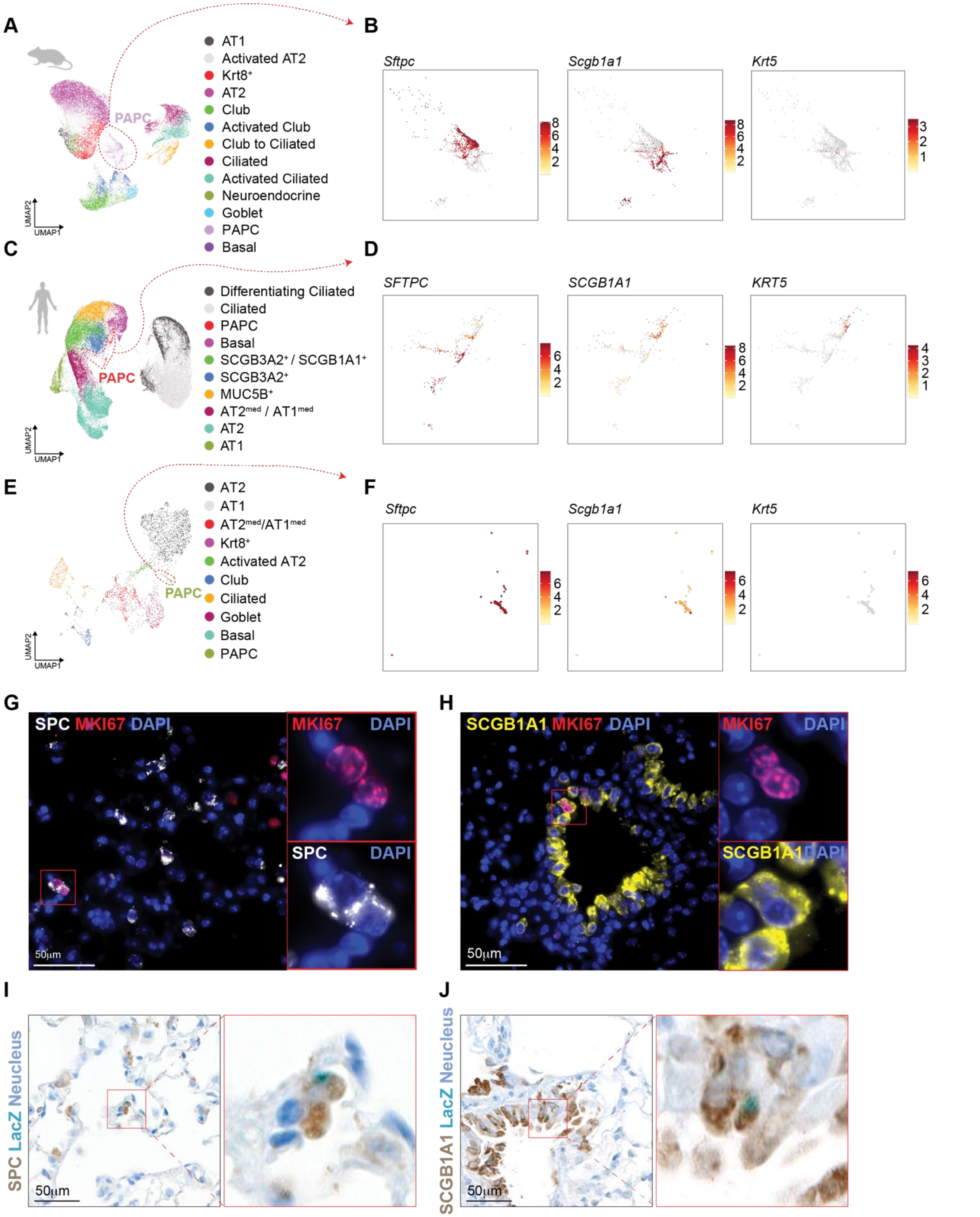


**Supplementary Fig. 9. PAPCs express various lung epithelial cell markers.**
**A** UMAPs displaying each cell cluster, colored by cell types of mouse lung shown in Figure 1A. **B** Feature plots displaying the expression of *Sftpc, Scgb1a1,* and *Krt5* in mouse PAPC cells. **C** UMAPs displaying each cell cluster, colored by cell types of human lung shown in Figure 1D. **D** Feature plots displaying the expression of *SFTPC, SCGB1A1,* and *KRT5* in human PAPC cells. **E** UMAPs displaying each cell cluster,

colored by cell types of mouse lung (*Pclaf* WT and KO at 7 dpi) shown in Figure 3B. **F** Feature plots displaying the expression of *Sftpc, Scgb1a1,* and *Krt5* in mouse PAPC cells. **G** Representative images of immunostaining for SPC and MKI67 using lung slides at 7 dpi shown in Figure 2. **H** Representative images of immunostaining for SCGB1A1 and MKI67 using lung slides at 7 dpi shown in Figure 2. **I** and **J** Representative images of immunostaining for SPC (**I**) and SCGB1A1 (**J**) in combination with X-gal staining, using lung slides at 7 dpi are shown in Supplementary Figure 2.

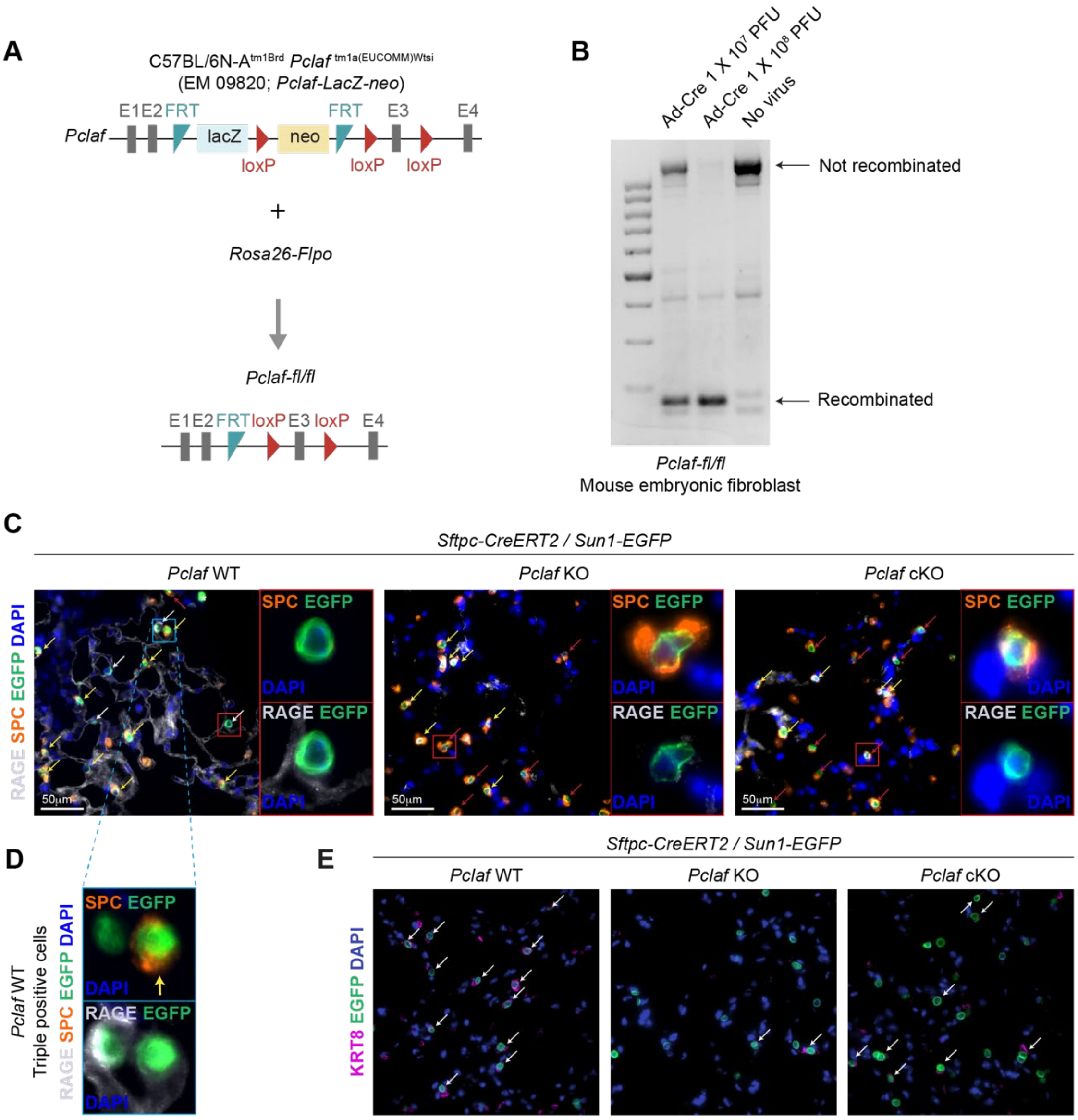


**Supplementary Fig. 10. Both germline and conditional KO of *Pclaf* inhibit AT1 cell generation from AT2 cells**

**A** Experimental scheme of generating of *Pclaf-fl/fl* mice. **B** Mouse embryonic fibroblasts (MEFs) were isolated from *Pclaf-fl (floxed)/fl* mice. *Pclaf-fl/fl* MEFs were treated with Ad-Cre viruses ($1 \times 10^7$ or $1 \times 10^8$ PFU). Conditional knock-out (cKO) by Ad-Cre was checked by PCR-based genotyping. **C-F** *Sftpc-CreERT2*; *LSL-Sun1-EGFP*; *Pclaf* WT, *Sftpc-CreERT2*; *LSL-Sun1-EGFP*; *Pclaf* KO, and *Sftpc-CreERT2*; *LSL-Sun1-EGFP*; *Pclaf-fl/fl* mice were instilled with bleomycin and then injected with tamoxifen for 5 consecutive days. At 14 dpi, lung tissues were collected and analyzed. Representative images of lungs; immunostaining for RAGE, SPC, and EGFP (**C**). White arrow; RAGE+/EGFP+ cell. Red arrow; SPC+/EGFP+ cell. Yellow arrow; RAGE+/SPC+/EGFP+ cell. Enlarged image of RAGE and SPC double-positive cell (**D**). Representative images of lungs; immunostaining for KRT8 and EGFP. White arrow; KRT8+/EGFP+ cell (**E**).

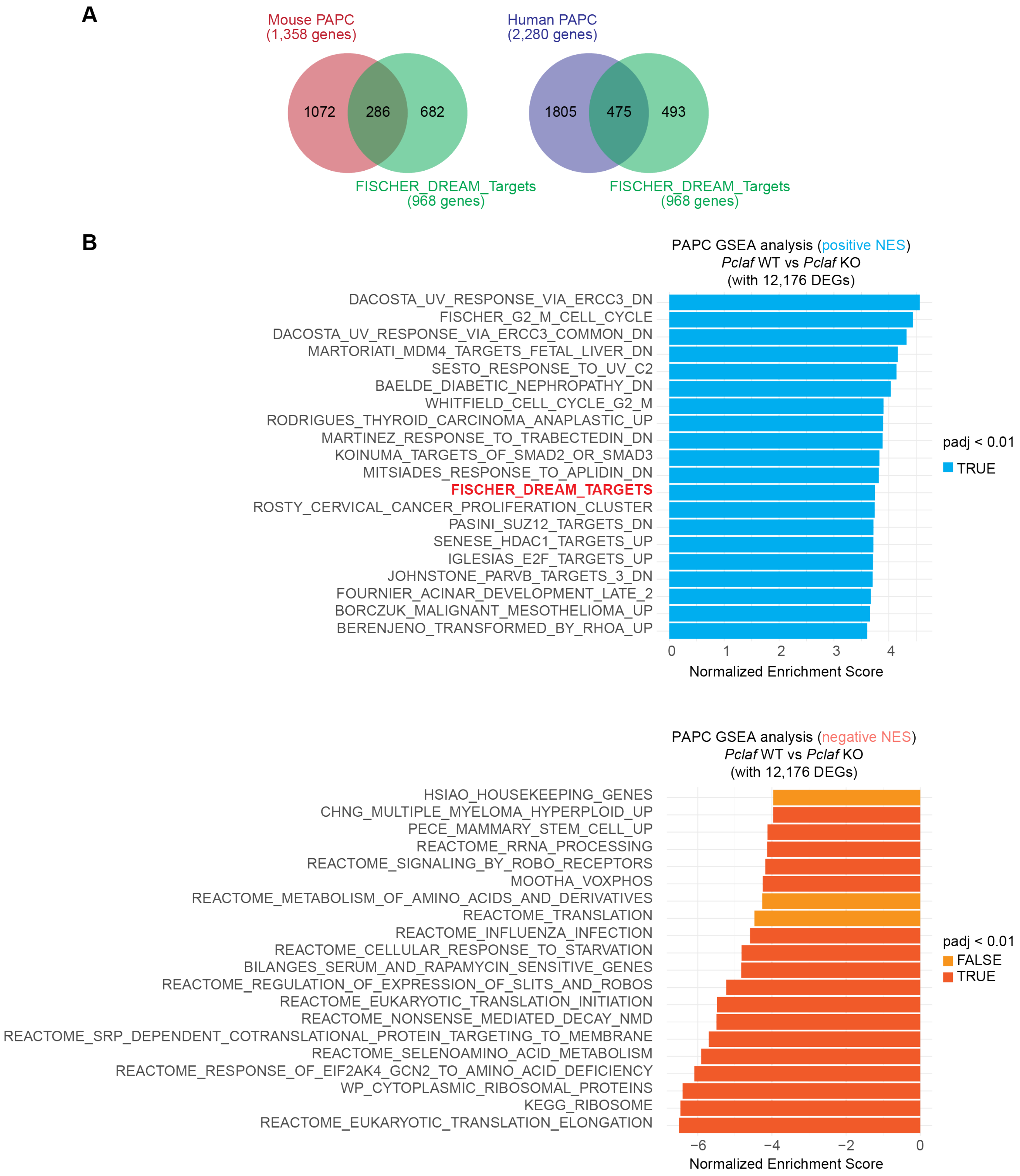


**Supplementary Fig. 11. *Pclaf* KO downregulates DREAM-target gene expression in PAPC.**
**A** Venn diagram analysis of DREAM target genes *vs.* genes specifically expressed in the PAPC clusters (mouse scRNA-seq dataset shown in Figure 1A or the human scRNA-seq dataset shown in Figure 1D). **B** Top 20 gene sets (*upper* panel) and bottom 20 gene sets (*bottom* panel) of GSEA with 12,176 DEGs between *Pclaf* WT PAPCs and *Pclaf* KO PAPCs in the scRNA-seq dataset shown in Figure 3.

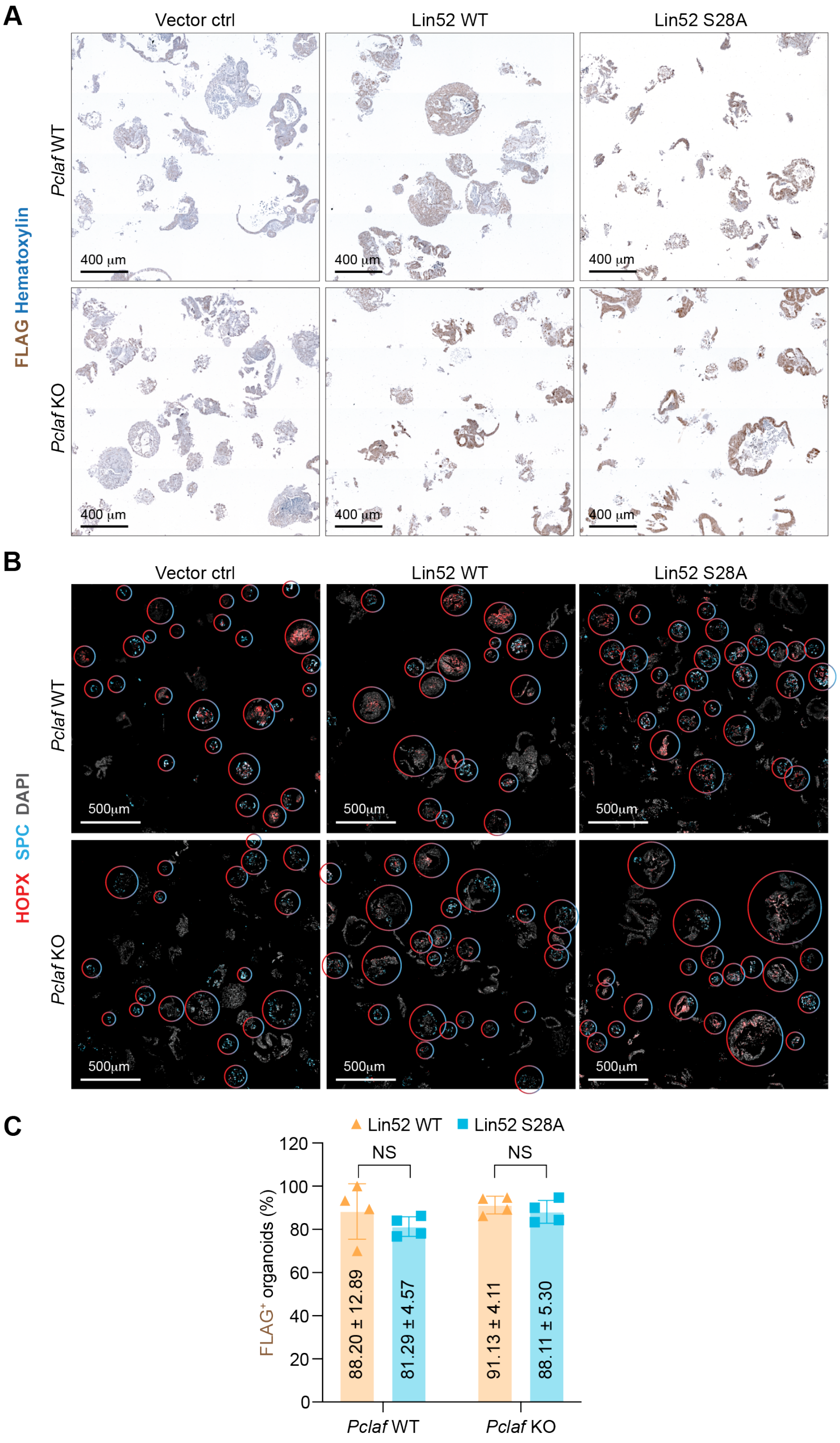


**Supplementary Fig. 12. Transduction efficiency of lentiviruses encoding Lin52-WT or Lin52 S28A in LOs** Isolated lung epithelial cells were transduced with RFP, Lin52 WT, or Lin52 S28A by lentivirus and then cultured with LO. **A** Representative images of chemically immunostained for FLAG. **B** Representative images of immunofluorescent (IF) staining for HOPX (AT1) and SPC (AT2) on day 14. The circles (in red-light blue mixed) indicate the relative ratio of $HOPX^{+}$ to $SPC^{+}$ cells in LOs. **C** Quantification of $FLAG^{+}$ LOs. The represented images and data are shown (n≥3); error bars: SD.

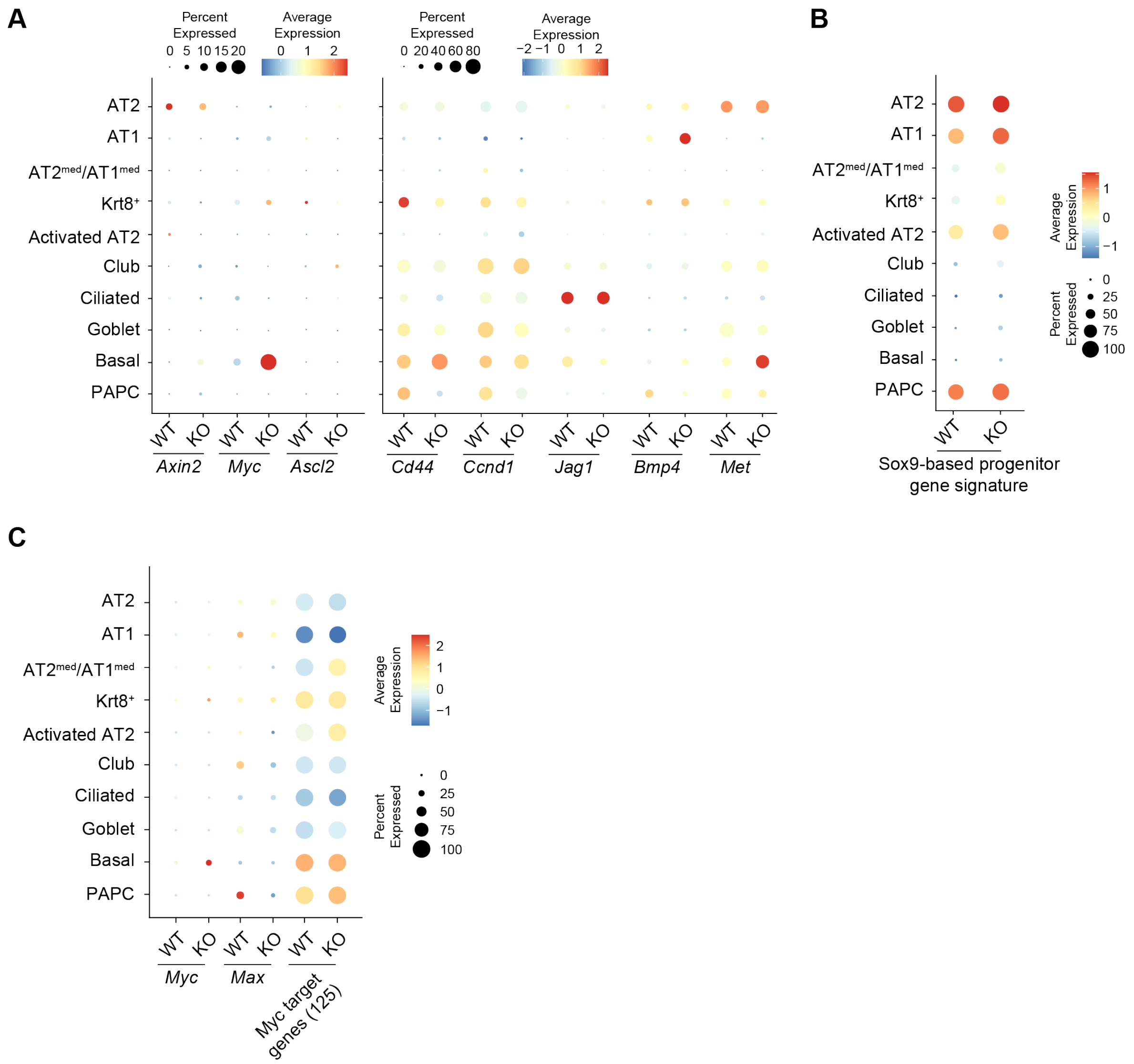


**Supplementary Fig. 13. Impact of *Pclaf* KO on Wnt signaling, Sox9, and Myc transcriptional signatures**
**A-C** Dot plots displaying each gene expression from the scRNA-seq dataset shown in Figure 3. Dot plots showing the expression of Wnt signaling target genes in each cell type (**A**). Dot plots depicting transcriptional module scores of the Sox9-based progenitor cell signature gene set (**B**). Dot plots showing the expression of *Myc* and *Max* and transcriptional module scores of the Myc target gene set (**C**). *Pclaf* KO barely alters Wnt signaling and Sox9-based progenitor transcriptional signature.

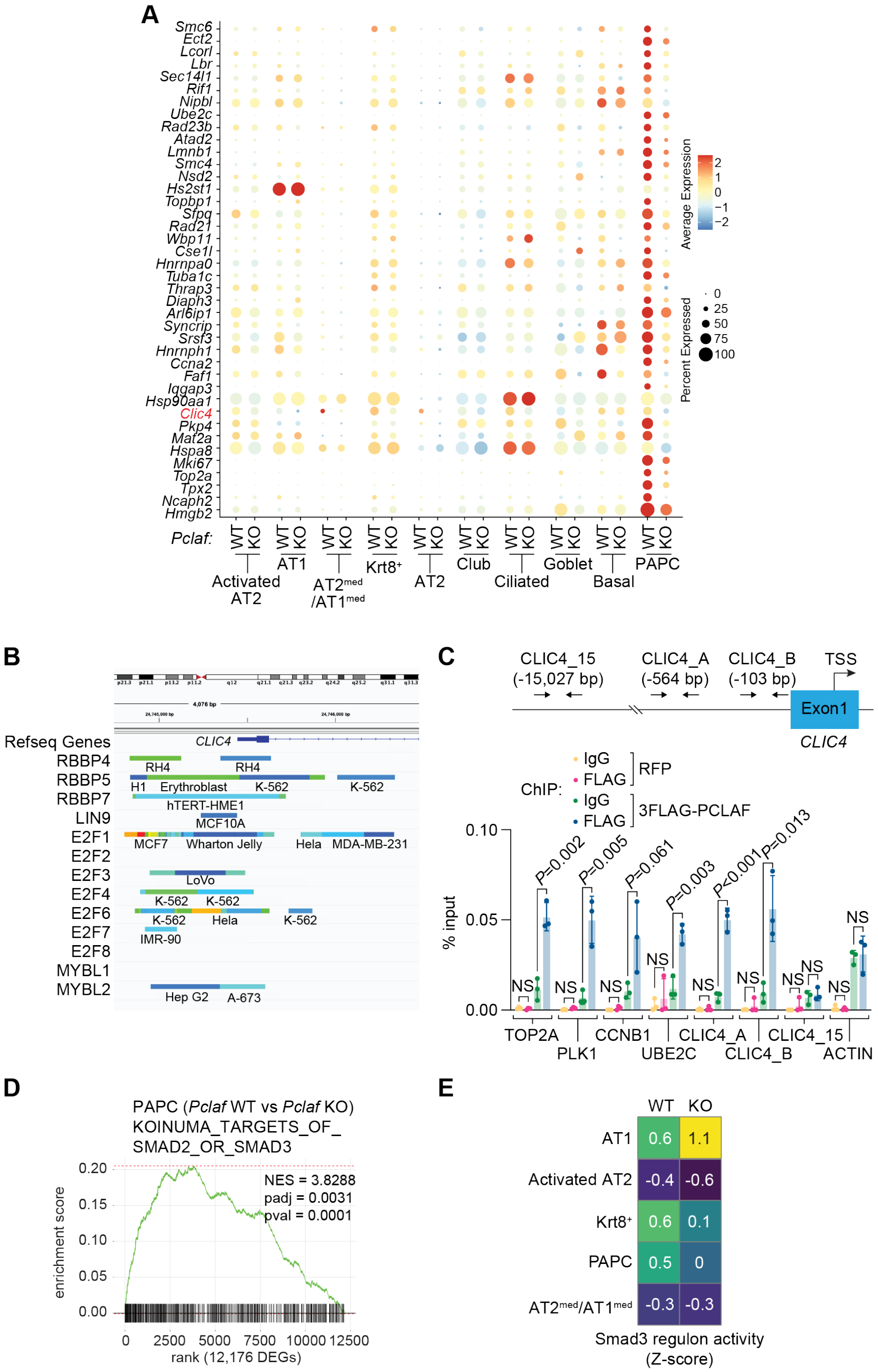


**Supplementary Fig. 14. *Clic4* downregulation is associated with decreased activity of TGF-β signaling in *Pclaf* KO PAPCs.**

**A** Dot plots displaying the top 40 differently expressed DREAM target genes in each cell type. **B** The DREAM complex components bind to the *CLIC4* promoter in ChIP-Atlas database (chip-atlas.org). **C** qPCR analysis using indicated primer sets targeting proximal promoter of DREAM target genes, *CLIC4*, or *ACTB*. ChIP was performed using anti-FLAG antibody. H358 cells ectopically expressing 3FLAG-*PCLAF* were used for ChIP. **D** GSEA of *Pclaf* WT vs. *Pclaf* KO in the PAPC cluster using the data set shown in Figure 3. The enrichment plot presents the gene sets of SMAD2 or SMAD3 target genes. **E** Z-score of Smad3 regulon activity by cell type and genome, analyzed by pySCENIC using the data set shown in Figure 3.

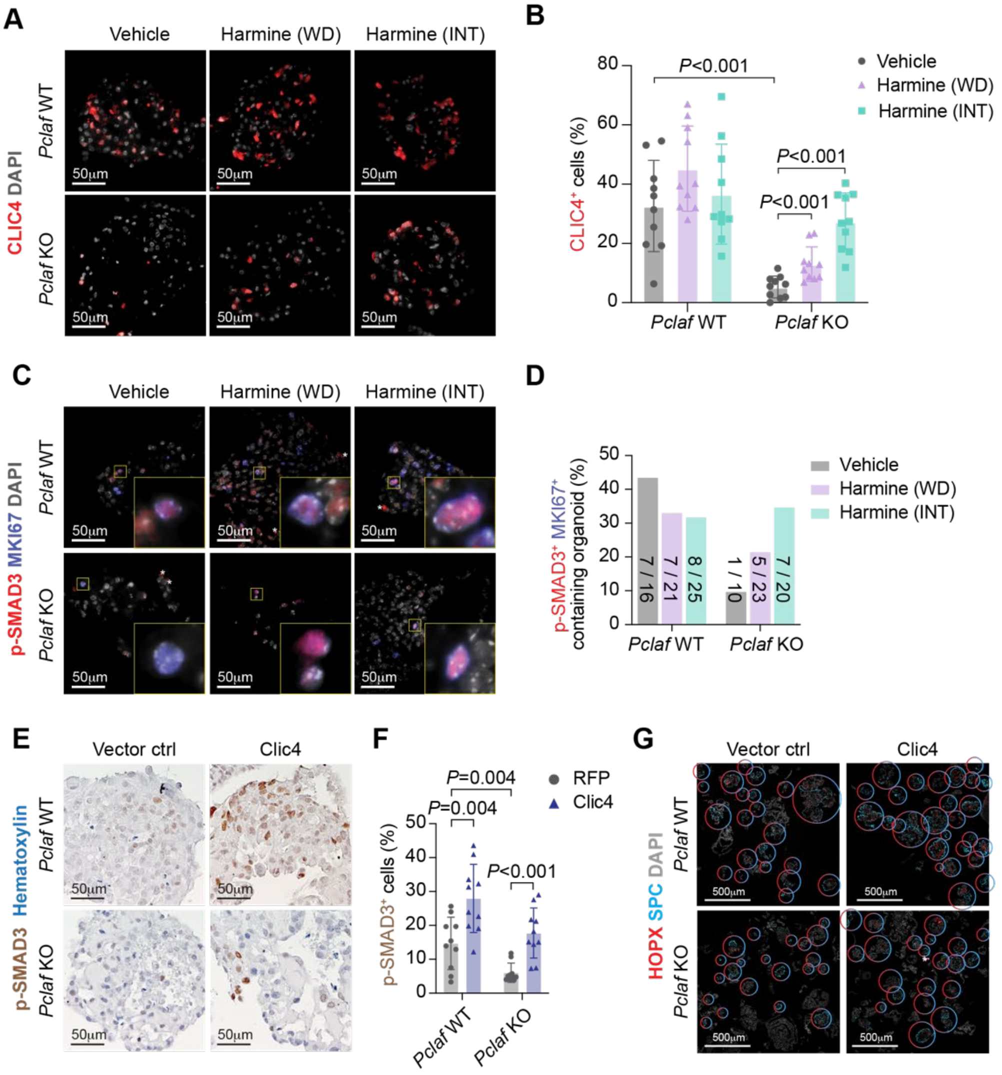


**Supplementary Fig. 15. Pharmacological or genetic activation of the DREAM axis rescues *Pclaf* KO-inhibited TGF-β signaling.**

**A**-**D** LOs treated with harmine (shown in Figure 4E) were immunostained with indicated antibodies. Representative images of IF staining for CLIC4 (**A**). Quantification of the CLIC4$^{+}$ cells (**B**). Representative images of IF staining for p-SMAD3 and MKI67 (**C**). Quantification of LOs containing p-SMAD3 and MKI67 double-positive cells (**D**). **E**-**G** Isolated lung epithelial cells were transduced with *RFP*- or *Clic4*-expressing lentiviruses and cultured with LO. Representative images of *Pclaf* WT and *Pclaf* KO LOs at day 14, immunostained for p-SMAD3 (**E**). Quantification graph of p-SMAD3$^{+}$ cells (**F**). Images of IF staining for HOPX (AT1) and SPC (AT2) on day 14. The circle color indicates the relative ratio of HOPX$^{+}$ to SPC$^{+}$ cells (**G**). Two-tailed Student's *t*-test; error bars: SD. The representative images are shown (n≥3).

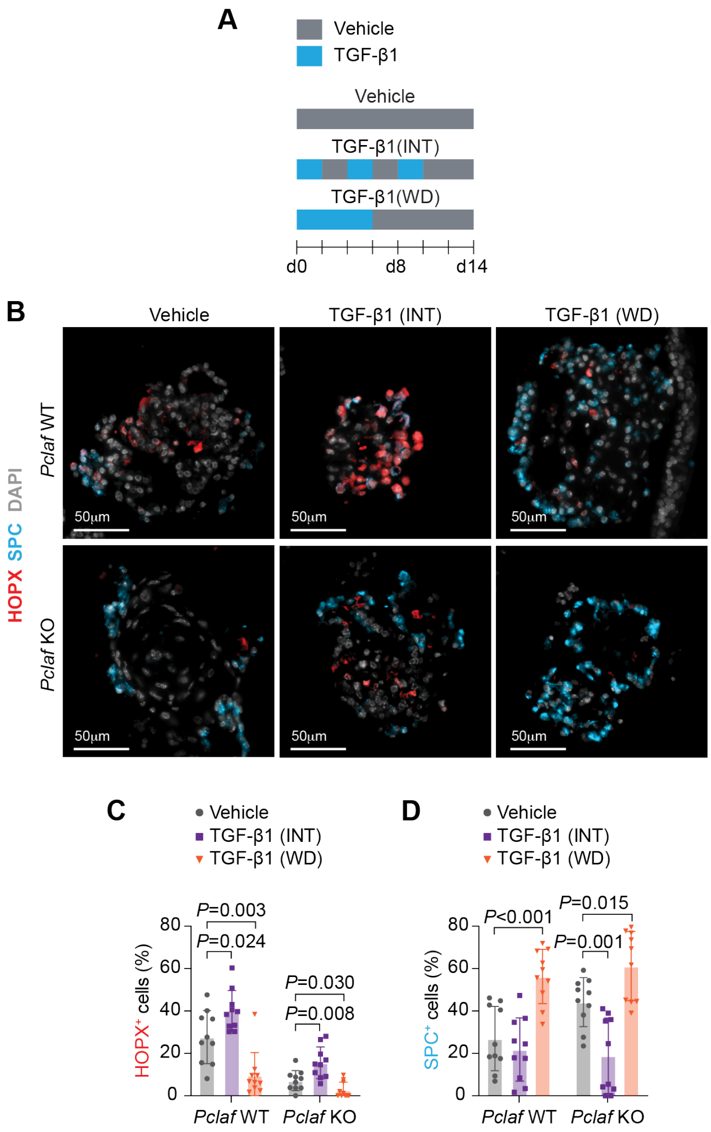


**Supplementary Fig. 16. TGF-β rescues *Pclaf* KO-impaired cell plasticity.**
Murine lung epithelial cells were cultured for LOs under stimuli of TGF-β1 (2 ng/ml). TGF-β1 was used to treat LOs for the first 6 days and withdrawn (WD). Alternatively, LOs were cultured with TGF-β1 intermittently (INT) at the indicated time points. **A** Experimental scheme for LO culture. **B** Representative images of IF staining for HOPX (AT1) and SPC (AT2) on day 14. **C**, **D** Quantification of $HOPX^+$ (**C**) and $SPC^+$ cells (**D**). Two-tailed Student's *t*-test; error bars: SD. The representative images are shown (n≥3). TGF-β1 (INT) rescued *Pclaf* KO-impaired alveolar cell plasticity (AT1 cell generation from AT2 cells). In contrast, TGF-β1 (WD) inhibited AT1 cell generation regardless of *Pclaf* WT and *Pclaf* KO. These data suggest that the temporal activity of TGF-β signaling is pivotal for AT2-to-AT1 cell transition. Of note, this experiment was designated by two rationales. First, signaling pathways elicit various outcomes depending on the spatiotemporal and dosage of signaling cues [3-5]. For example, intermittent or continuous stimulation of parathyroid hormone (PTH) can result in either increased or decreased bone mass, respectively [6]. Second, the balance between proliferation and differentiation is critical in organoid culture in general and TGF-β signaling has been shown to inhibit the proliferation of alveolar cells [7]. Since TGF-β generally inhibits cell proliferation, we thought that stimulation of TGF-β during the whole period would severely inhibit organoid formation and growth. Thus, to reduce the inhibitory effect of TGF-β, we cultured the LOs under TGF-β stimulation of half period of culture with two different models. The INT model aimed to continuously alter the organoid culture condition between TGF-β1 stimulation and depletion, while the WD model aimed to provide continuous TGF-β1 stimulation at the early stage of LO culture.

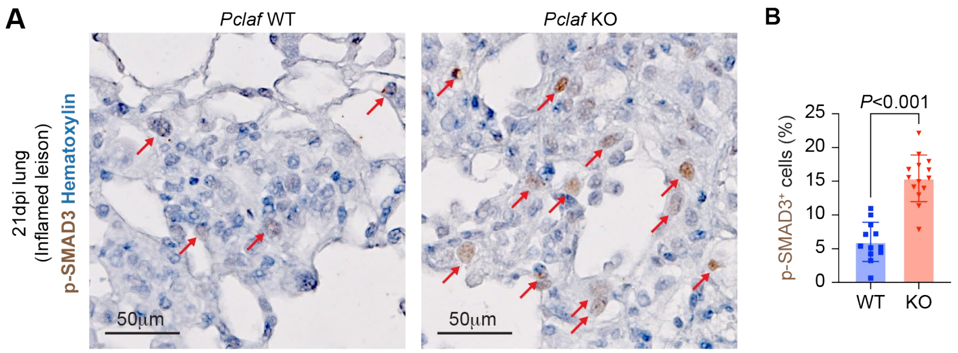


**Supplementary Fig. 17. Elevated TGF-β signaling in inflamed lesions of *Pclaf* KO lung tissues**

**A** Images of immunostaining for p-SMAD3 of the inflamed lesion from *Pclaf* WT and *Pclaf* KO lung tissues at 21 dpi. **B** Quantification graph of p-SMAD3$^+$ cells. Two-tailed Student's *t*-test; error bars: SD. The representative images are shown (n≥3).

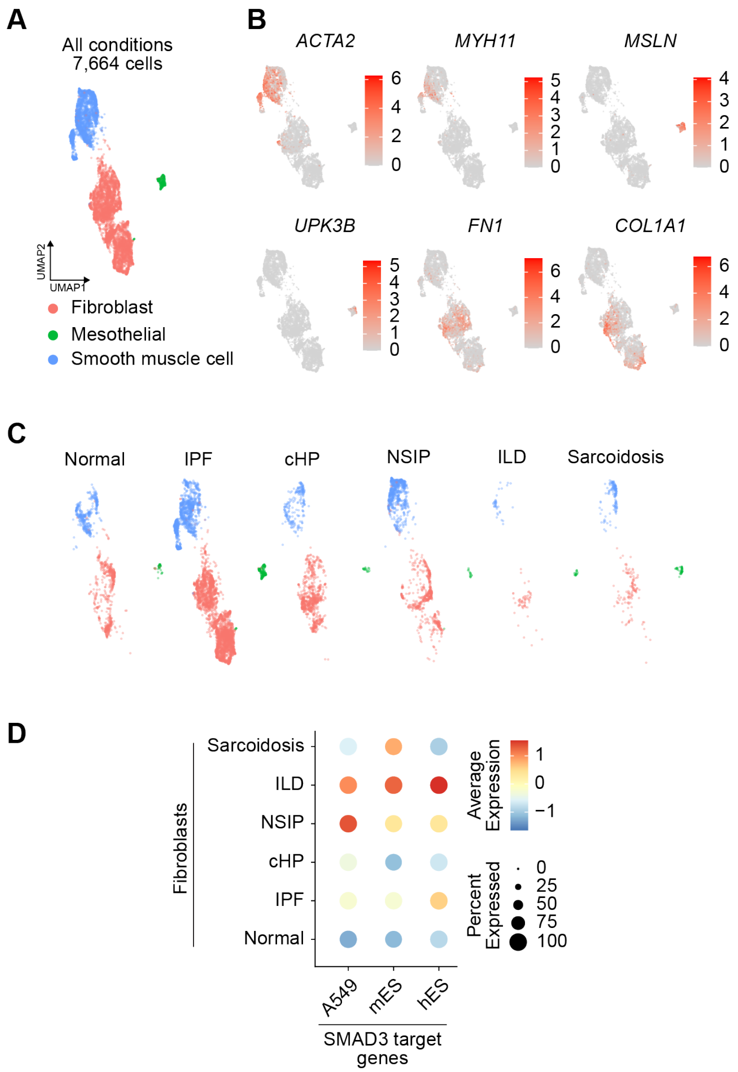


**Supplementary Fig. 18. Elevated TGF-β signaling in lung fibroblasts of IPF patients**

**A** UMAP showing the Mesenchymal compartment (*EPCAM*$^-$, *PTPRC*$^-$, and *PECAM*$^-$ cells) from the human scRNA-seq dataset (GSE135893; normal, IPF, cHP, NSIP, ILD, and Sarcoidosis). **B** Feature plots of indicated marker genes: *ACTA2* and *MYH11* for smooth muscle cells; *MSLN* and *UPK3B* for mesothelial cells; *FN1* and *COL1A1* for fibroblasts. **C** UMAP embedding displays cells colored by cell types split by each disease type. **D** Dot plots showing the expression of indicated genes and module scores of SMAD3-target gene sets in human lung fibroblasts.

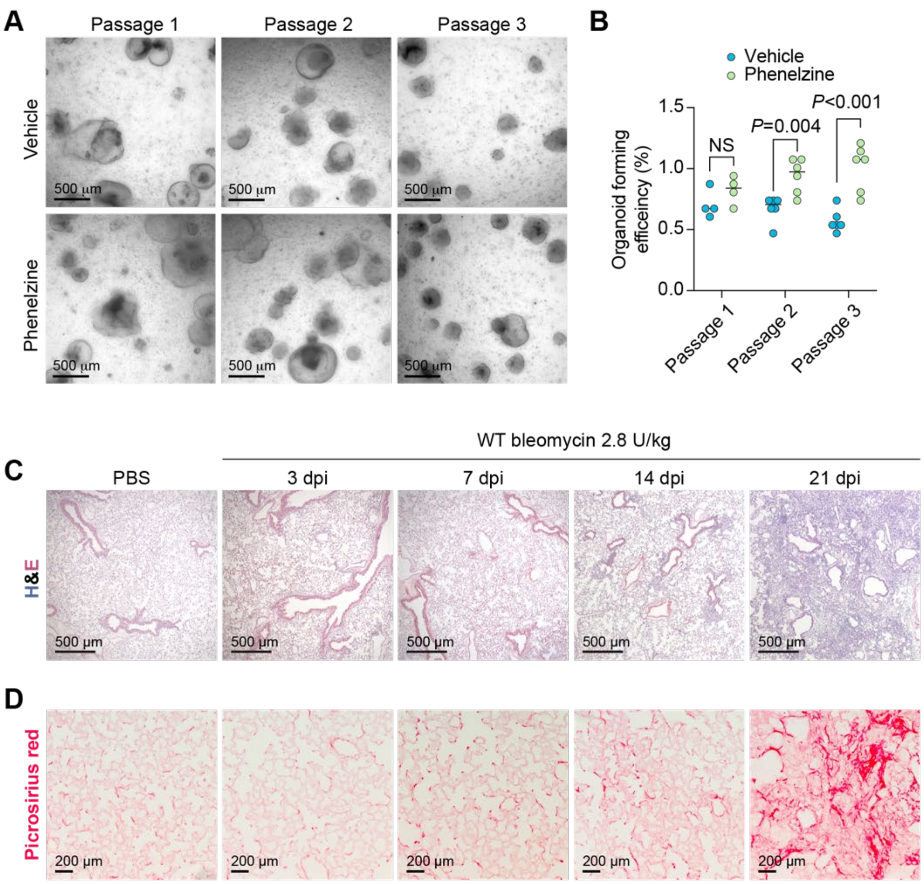


**Supplementary Fig. 19. Phenelzine promotes LO formation.**

**A** Images of LOs with phenelzine (10 μM) at 12 days of indicated passage. **B** Quantification graph of lung OFE at 12 days of passage. **C, D** 8-week-old C57BL/6 mice were treated with PBS (n=3) or bleomycin (2.8 U/kg, n=3 of each group) by intratracheal instillation. Lung tissues were collected at indicated time points. Lungs with PBS were collected at the same time point of 21 dpi. Representative images of H&E staining at the indicated time points (**C**). Representative images of picrosirius red staining at the indicated time points (**D**). Two-tailed Student's *t*-test; error bars: SD. The representative images are shown (n≥3).
